# Global mismatches between crop distributions and climate suitability

**DOI:** 10.1101/2021.04.15.439966

**Authors:** Lucie Mahaut, Samuel Pironon, Jean-Yves Barnagaud, François Bretagnolle, Colin K. Khoury, Zia Mehrabi, Ruben Milla, Charlotte Phillips, Delphine Renard, Loren H. Rieseberg, Cyrille Violle

## Abstract

The selection of new crops and the migration of crop areas are two key strategies for agriculture to cope with climate change and ensure food security in the coming years. However, both rely on the assumption that climate is a major factor determining crop distributions worldwide. Here, we show that the current global distributions of nine of twelve major crops strongly diverge from their modelled climatic suitability for yields, after controlling for technology, agricultural management and soil conditions. Comparing the climatic niches of crops and their wild progenitors reveals that climate suitability is higher outside the native climatic range for six of these nine crops while all of them are farmed predominantly in their native ranges. These results show that agricultural strategies coping with climate change will be unsuccessful unless they fully consider the social, cultural, and ecological factors underpinning crop distributions.

---

Ensuring food security while adapting to climate change, and preserving the environment, is a central challenge for humanity^1,2^. Many solutions have been proposed, including the migration of crop areas^3–5^ and the cultivation of novel species and varieties^6^ to identify the most promising species and regions for food production under different scenarios of climate change. Most studies on agricultural adaptation to climate change consider that climate is a major factor determining the growth and productivity of crops, and that crop distributions can be optimized for climate by human societies. However, such proposals often do not account for multiple agricultural management (e.g. irrigation), cultural (e.g. preferences for certain crops from farmers and consumers^7^), socio-economic (e.g. agricultural policies, subsidies, markets, and international trade^8,9^) and other ecological (e.g. pest pressure^10^) factors, that jointly determine the current biogeographic patterns of crops.

The extent to which non-climatic factors matter on a global level for climate adaptation in agriculture can be assessed through identifying mismatches between current climate suitability and crop distributions. Analysing crop distribution-climate suitability (mis)match is thus critically important for any discussion on global climate adaptation in agriculture and more generally, for understanding how humans and crops have co-evolved with climates.

Here we investigate the relationship between the global distributions of twelve major food crops (Extended Data Table 1) and their climate suitability, using a global database of crop distributions^11^ and a crop-specific climatic niche model based on mean annual temperature (MAT) and total annual precipitation (TAP). Both MAT and TAP are essential to the survival and growth of domesticated and wild plant species and have recently been related to the global distribution of croplands^12^. Within the two-dimensional climatic space of each crop, we assessed climate suitability by predicting the effects of MAT and TAP on crop yield while controlling for agricultural inputs (i.e. irrigation and fertilization) and socio-economic factors (i.e. gross net product and human development index) to account for differences in terms of technological inputs, as well as numerous biophysical factors, including soil conditions and topography (see Methods). We then tested for a significant correlation between climate suitability and the fraction of cropland allocated to each crop (hereafter *crop area*), and mapped the global distribution of the (mis)match between crop area and climate suitability (see Methods). Further, we investigated the role of crop origins and agricultural expansion in the relationship between the global distribution of crops and climate suitability. We used native occurrences of the wild progenitors of each crop (Extended Data Table 1) to define their *native* versus *expanded* climatic ranges (i.e. portion of the crop climatic space occupied by its wild progenitors and by the crop only, respectively) and we compared crop area and climate suitability between the two ranges.

The optimal climatic conditions for yield (Figure 1) and the distribution of cultivated areas within crop climatic spaces (Figure 2) vary widely from one crop to another. Overall, our analysis shows that the current global distributions of individual crops poorly match their climatic potential. Among twelve major crops, the cultivation of only three - groundnut, maize and soybean - predominates in climates that are predicted to be the most suitable for their yields (blue in Figure 3). Conversely, cassava, rapeseed, rice and sunflower are mostly cultivated in climates of low suitability for their yields (red in Figure 3). Moreover, the fraction of cropland allocated to barley, potato, sorghum, sugarbeet and wheat is not significantly correlated with their predicted climatic suitability (black in Figure 3). Adding the seasonality of temperature and precipitation (i.e. two potentially important climatic factors for yields) to crops’ climate suitability models (see Methods) confirms that climate suitability is generally a poor predictor of realized crop distributions (Extended Data Figure 1). The main differences with the calculation of climate suitability based on MAT and TAP only are observed for wheat, whose distribution of cultivated areas becomes negatively correlated with the suitability of climate, and maize and soybean, whose area-suitability correlations become non-significant (Extended Data Figure 1).

**Figure 1:**
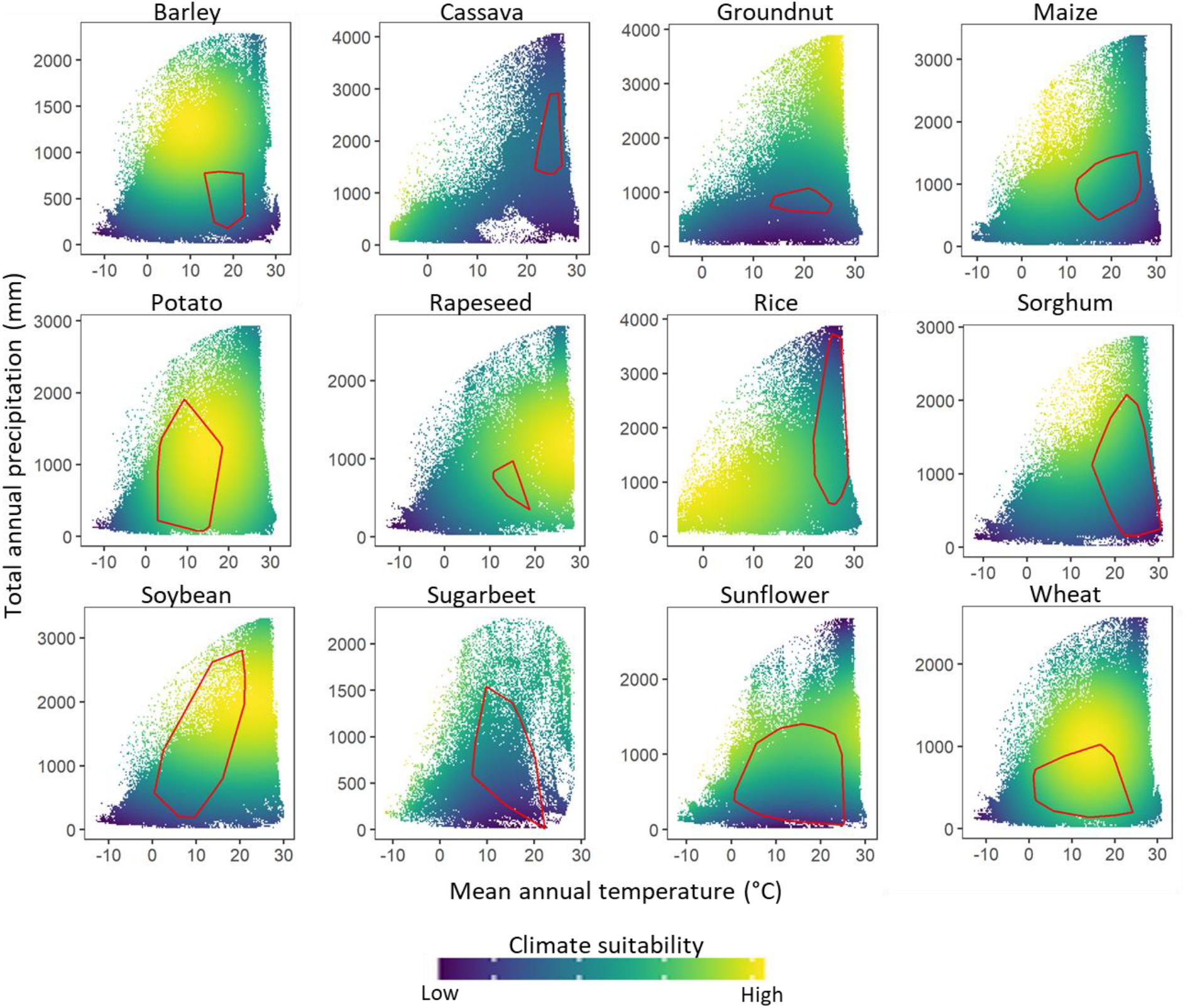
Climate suitability for the production of 12 major crops at the global scale. Climate suitability is predicted for each crop by modelling the effects of mean annual temperature (MAT) and total annual precipitation (TAP) on yield while controlling for the effects of various agricultural, socio-economical soil and topographic factors (see Methods). Red polygons delineate the climatic space occupied by crop wild progenitors (i.e. native climatic range).

**Figure 2:**
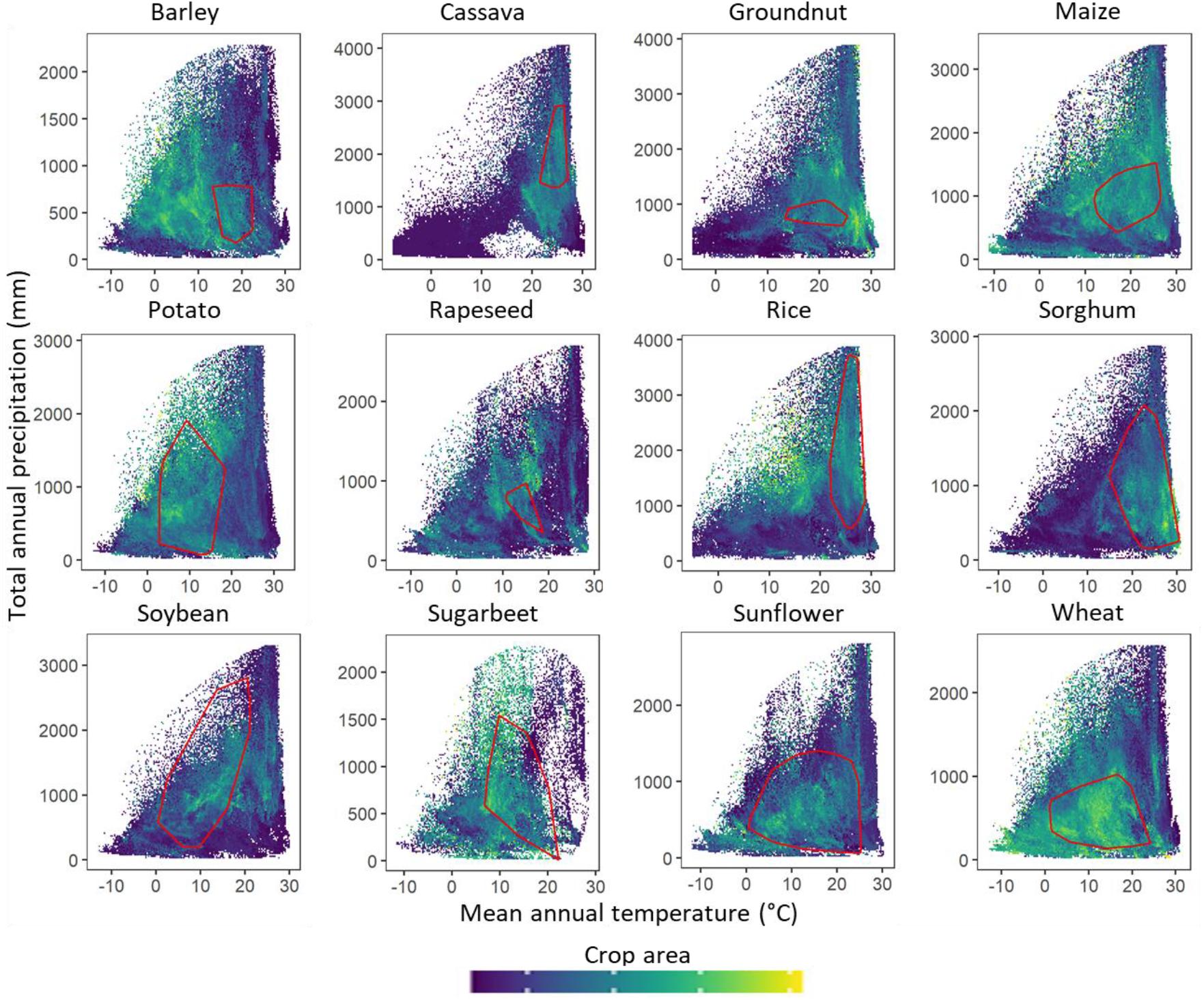
Distribution of the areas of 12 major crops within their two-dimensional climatic space. Crop area correspond to the fraction of total, available cropland devoted to each crop. Red polygons delineate the climatic space occupied by crop wild progenitors.

**Figure 3:**
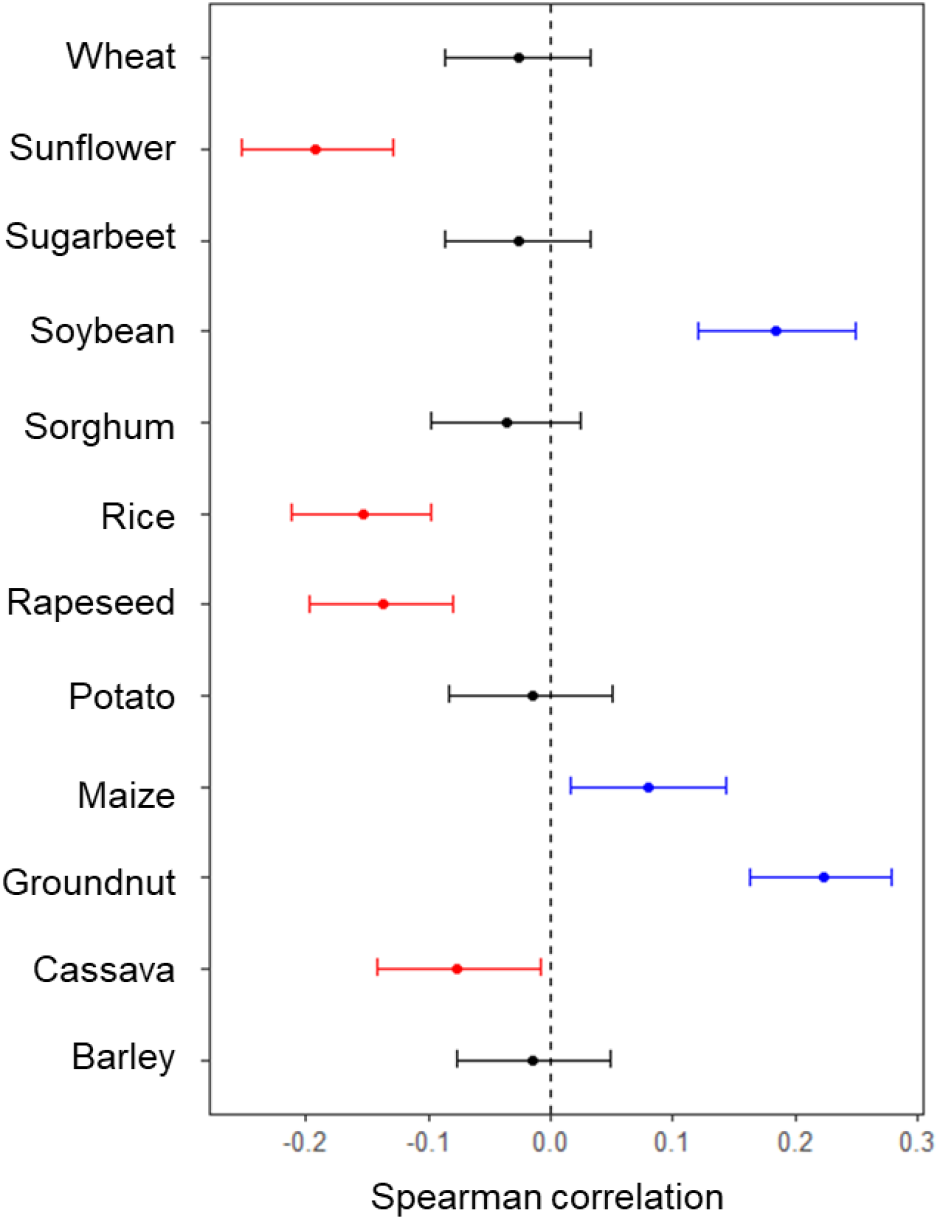
Correlation between the fraction of available cropland devoted to each crop and climate suitability. Horizontal bars show confidence intervals (alpha = 0.05) computed from 1000 random resamplings (see Methods). The correlation is considered significant if the confidence interval does not include 0. Black: non significant correlation; Blue: positive correlation; Red: negative correlation.

The strong mismatch between modelled climatically suitable cropland areas and current cultivation can indicate local limits to crop production, including climate, but can also allude to the importance of many other ecological or socio-cultural drivers (Figure 4). Geographic zones where the allocation of land to a crop is lower than predicted by its climatic suitability represent 78-98% of the global crop area (positive mismatch scores; blue areas in Figure 4). Different causes can explain such a dominant pattern, including the fact that some crops share similar climatic optima (e.g. maize and sorghum; wheat and potato; rapeseed and soybean, Figure 1), leading to competition for space. In addition, local food systems, and especially smallholder ones, typically cultivate a diversity of crops to serve multiple purposes (livelihood, subsistence, pest control^13,14^). Similarly, management factors such as multiple cropping systems^15^ and year-to-year crop rotations^16^ can create patterns of crop distributions that deviate from the modelled climate optima for single crops. The presence of natural enemies (pests and pathogens) may be another important explanation for the apparent mismatch between crop distributions and their climate optima for yield. Indeed, crop pests tend to share the same climatic niche as their host^10^ and thus farmers may choose to reduce the area allocated to a crop that is too intensely exposed to pests. We test this hypothesis for sunflower, using occurrence records of *Sclerotinia sclerotiorum* (adapted from Mehrabi et al. 2019 ^12^), a major sunflower pathogenic fungus worldwide. We find that the mismatch between climate suitability and the fraction of cropland allocated to sunflower is lower in pest free areas than in areas where the pest occurs (Extended Data Figure 2), suggesting a strong ecological anchoring of the mismatches between climate suitability and current cultivation areas.

**Figure 4:**
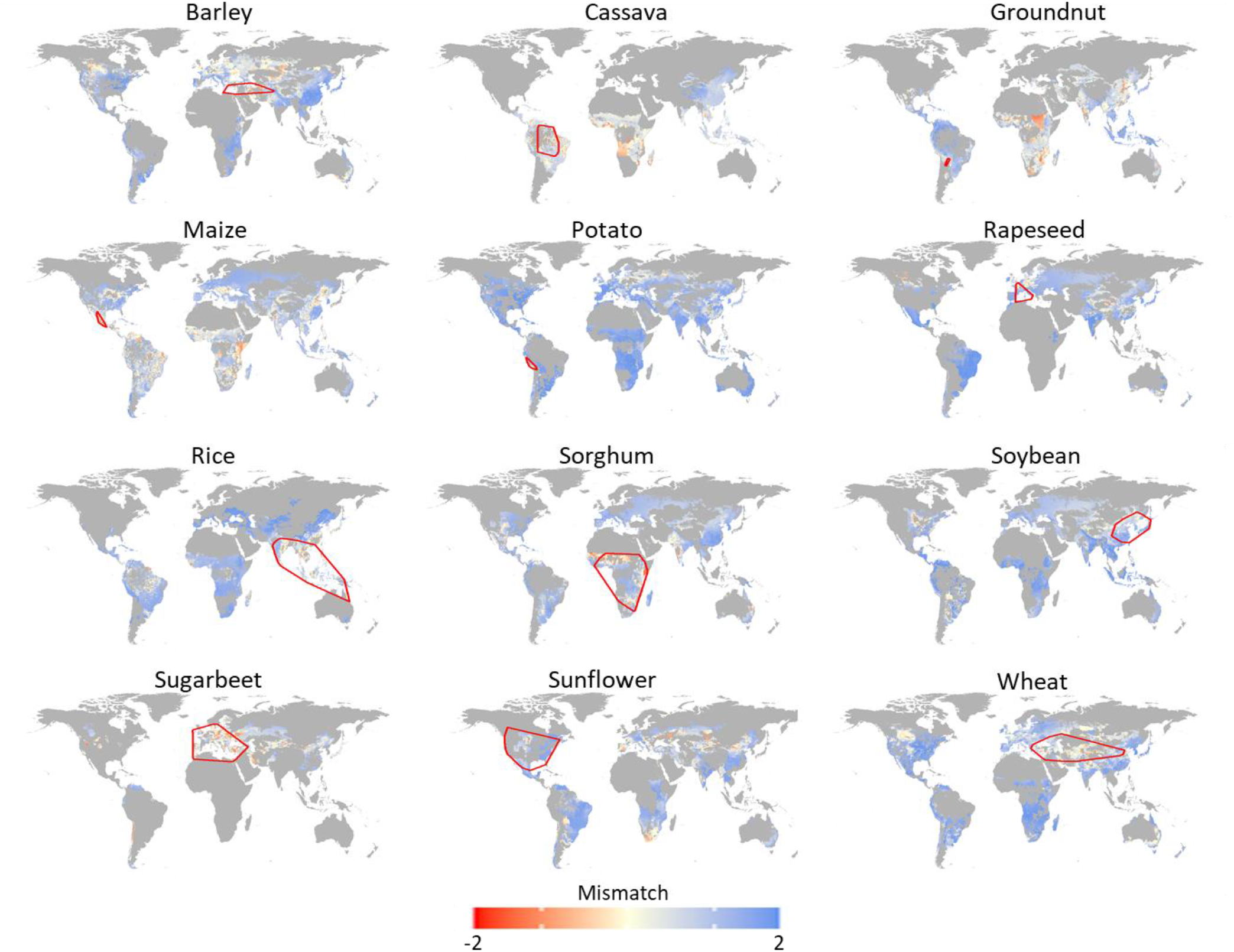
Mismatches between the fraction of available cropland devoted to each crop and climate suitability for their yield. Mismatch is computed as the log ratio of climate suitability divided by the fraction of cropland each crop occupy (see Methods). Positive scores (blue) are regions where crop areas are low relative to climate suitability. Negative values (red) indicate regions where cultivated areas are large with respect to climate suitability. Yellow designates regions where cultivated areas and climate suitability are in equilibrium. Red polygons delineate the geographic space occupied by crop wild progenitors (i.e. native range).

Our findings also highlight regions where the fraction of cropland devoted to a crop is large relative to the suitability of climate for its yield (2-21% negative mismatch scores; red areas in Figure 4). This is particularly true in (1) Eastern Europe and Central Asia for barley, sugarbeet and sunflower; (2) Central and Eastern Africa as well as South America for cassava and maize; (3) Eastern and South Africa, South India and Eastern Asia for groundnut and (4) Western and Eastern Africa for sorghum (Figure 4). Deeper insights into the social systems in which these crops are grown may explain these patterns. For example, cassava is a key crop for food security across the African continent. It represents a lifeline when other crops fail due to droughts, pest outbreaks, especially during the hunger season^17^. Thus even if yields may be comparatively low, they are reliable under marginal conditions^17^. Other factors not related to yield or food security likely play out in other situations. For example, agricultural policies, and market distortions such as fixed prices and subsidies, can help explain why maize is cultivated to a wider spatial-climatic extent than its climate optimum might predict^9^.

Major crops by definition have undergone substantial geographical expansion from their respective regions of origin to the rest of the world^18^. They have adapted to new climates through historical globalization, including trade and innovations in agricultural practices, especially irrigation^19^. However, the resulting current distributions of crops in comparison to the native distributions of their wild progenitors, with regard to climate suitability, remains poorly quantified^18^. We find that the fraction of agricultural land allocated to a given crop is higher in native than in expanded climatic ranges for all crops except barley and groundnut (Figure 2 & Extended Data Figure 3). Since areas of early crop domestication likely occur in, or nearby, the native distributions of their progenitors^20,21^, our results reveal that major crops are farmed predominantly in the same climates as their geographic origins, potentially indicating the biological constraints of crop species to wide climate adaptation. Native climatic ranges are also more associated with climatic suitability than expanded ones for cassava, maize, potato, rapeseed, sorghum and wheat (Figure 1 & Extended Data Figure 3), suggesting that these crops are still better adapted to the climates of their origins. Conversely, native climatic ranges are less suitable for barley, groundnut, rice, soybean, sugarbeet and sunflower when compared to climate suitability in crops’ expanded ranges (Figure 1 & Extended Data Figure 3). These results corroborate suggestions that humans have introduced crops into climatically more suitable areas outside their native ranges^1^, although we cannot rule out the possibility that climate might also have changed since domestication into less suitable conditions in native ranges^22^. Perhaps most importantly, varietal selection, mutations, and plant breeding has modified the ecological requirements of crops so that they have become much more widely adapted^23^.

The relationship between wild species distributions and performance (usually in some yield equivalent, such as fitness) is a core issue in the ecological niche theory^24,25^. Our investigation of cultivated species shows that climate suitability is a poor predictor of the current distribution of crops, even after controlling for agricultural management, technology proxies and soil conditions. Building crop climatic niches has previously been proposed as an approach to ensure global food security by guiding species selection^6^ and by identifying new potential areas to cultivate^5,26,27^ under future climate scenarios. However, our work shows the importance of other, non-climatic factors, in determining crop distributions today. The mismatches we found between crop distribution and climate suitability could serve as a basis for further research to better understand the determinants of the global distribution of crops, notably the role of ecological factors such as the presence of pests and their predators, as well as key social factors, such as nutrition, resilience and markets. In turn, greater insights into these multiple factors could help to integrate the climatic requirements of crops into agricultural planning. Finally, our study tells a remarkable story of how humans and crops have co-evolved with climates over the course of agricultural history, well beyond climatic constraints generally considered to be highly determinant of species distributions. Extending our approach to less studied ‘orphan crops’^28,29^ which haven’t been moved around as much as major crops and which are less heavily managed will help to extend the insights provided here further.

## Materials and Methods

### Agricultural data set

We studied barley, cassava, groundnut, maize, potato, rapeseed, rice, sorghum, soybean, sugarbeet, sunflower and wheat (Extended Data Table 1). The global distributions of harvested areas and yields were extracted from public data sources^11^ in the form of rasters of 5 arc-minutes resolution (~10km). These 12 crops were chosen because (1) they are widely cultivated worldwide and provided more than 70% of food globally; (2) data on the amount of fertilizers – known as a major driver of crop yields globally - are available at a 5 arc-minutes resolution^30^ and (3) their wild progenitors can be identified in the literature^20^ and have at least 20 recorded occurrences in the wild (Extended Data Table 1). We used the global distribution of cropland^31^ to compute the fraction of available cropland allocated to each crop at a 5 arc-minutes resolution. Nutrient application on major crops and the percentage of land area equipped for irrigation^32^ were downloaded as 5 arc-minutes rasters.

### Crop progenitors

We identified 23 progenitors of the 12 crops using published literature^21^. Wheat includes *Triticum aestivum* and *Triticum durum*, which are the two main wheat species farmed worldwide. Therefore, we considered the progenitors of both wheat species. Rice also includes two different species, *Oryza sativa* (Asian rice) and *Oryza glaberrima* (African rice). However, because *O. sativa* is by far the principal cultivated rice species, we only considered the wild progenitors of the Asian rice in our analysis. We extracted progenitor occurrence records from five global and two regional databases. We used the *BIEN* and *rgbif* packages in R to download global occurrence records from the Botanical Information and Ecology Network^33^ and the Global Biodiversity Information Facility^34^ databases, respectively. In addition, we extracted global occurrences of crop wild progenitors from “A global database for the distributions of crop wild relatives”^35^, the BioTIME database^36^ and GENESYS^37^. We downloaded further regional occurrences using the RAINBIO database, which contains records for Sub-Saharan African vascular plants^38^ and *species*Link, a national database for plant and animal occurrences in Brazil^39^. Because occurrence data are often inaccurate^40^, we removed records with no coordinates and that were documented before 1950. We used the *CoordinateCleaner* package in R^41^ to remove occurrence records found within a 1 km radius of country and capital centroids, with equal longitude and latitude coordinates, or assigned to institutional locations such as botanical gardens, herbaria or the GBIF Headquarters. We also removed records with high coordinate uncertainty (over 10 km), cultivated records (e.g. breeding/research material, advanced/ improved cultivar, GMO and those found in markets or shops, institutes/ research stations and genebanks and from seed companies), and records located in the sea/oceans. For progenitors with the same species name as their crop, we only extracted records confirmed as wild and ignored any record with an unknown cultivation status. In addition, we removed records outside of the species reported native range to ensure no introduced and cultivated record was included. Native ranges were identified at administrative levels according to the USDA Agricultural Research Service, Germplasm Resources Information Network^42^. We removed duplicates by keeping only one record for each species per 20 km grid cell. After discarding the less accurate occurrence records, 4064 unique occurrence points remained for 23 wild progenitor species of the 12 crops (Extended Data Table 1).

### Climate data

We acquired mean annual temperature (MAT) and total annual precipitation (TAP) data at 30 arc-seconds (~1 km) for the years 1979–2013 from the CHELSA database^43^. We also used temperature and precipitation seasonality to account for their potential effects on crop yields.

### Soil, topographic and socio-economic data

We integrated soil pH, soil organic content, soil water capacity, slope as well as the human development index and gross net product, two proxies of technological inputs, to control for their effects on crop yields. These layers are freely accessible at the URLs provided in Extended Data Table 2.

### Climate suitability

For each crop, we estimated climate suitability by modelling the effects of MAT and TAP as second order polynomials on crop yield while controlling for the effects of fertilisation and irrigation (as second order polynomials) as well as all other soil, topographic and socio-economic factors. We upscaled all the agricultural (i.e. crop yields and areas), climatic, soil, topographic and socio-economic layers to 10 arc minutes resolution (~20 km). We worked at such a resolution because climate is expected to be the main driver of species distribution at large scales, whereas other factors might become more important with lower grain size^44^. In doing so, we also minimized incorrect assignment of climatic variables to the occurrence records for which precision is not always communicated by data sources. We used the estimated coefficients of this model (Extended Data Table 3) to compute climate suitability as the yields predicted only by MAT and TAP. Crop yields were square root-transformed to assure linearity. We then tested the robustness of the climate suitability model on 1,000 data subsets, each constructed by randomly sampling 1,000 points (without replacement and without stratification) from the data set (pixels) of each crop. We identified covariates that have a significant effect on crop yields using the confidence intervals (alpha = 0.05) of the estimates from these random re-samplings. In addition, we used the sums of squares to quantify the amount of global variability in crop yields explained by the different factors (Extended Data Table 3). Finally, we repeated this procedure by adding temperature and precipitation seasonality to the climate suitability model (Extended Data Table 4).

### Correlation between crop areas and climate suitability

We calculated the Spearman correlation index between the fraction of available cropland allocated to each crop and climate suitability to test whether the global distribution of crops matches their climate suitability. Because sample sizes were extremely large (i.e. between 52 011 and 176 318 pixels; Extended Data Table 1), we randomly sampled 1000 pixels 1000 times and computed the confidence interval of the Spearman correlation indices to test for their significance (alpha = 0.05).

### Mismatch index

To compare the spatial distribution of climate suitability (Extended data Figure 4) and crop areas (Extended data Figure 5), we computed for each crop a mismatch index as the log-ratio of climate suitability divided by the fraction of cropland covered by each crop in each 10 arc-minutes pixel. We previously scaled both variables between 1 and 2 so that mismatch values vary between - 2 (climate suitability is low with respect to cropland proportion) and +2 (climate suitability is high relative to cropland proportion; Figure 4).

### Native and expanded climatic ranges

We defined a crop’s native climatic range by drawing a convex polygon around the occurrence points of the pool of its wild progenitors^6^ within the two-dimensional climatic space of each crop. By contrast, crop expanded range refers to the part of this two-dimensional climatic space that is not covered by the progenitors. Although several techniques can quantify species’ climatic range^45^, convex hull methods appear to be particularly relevant for large-scale agricultural applications and its relatively simple concept makes it easily interpretable^6^. Because the use of convex polygons is sensitive to outliers, we removed occurrence points with Mahalanobis distance values ≥ 10 from the wild progenitors’ niche space defined by MAT and TAP^6^. This threshold was selected based on visual inspection of the data.

### Difference in crop areas and climate suitability between current and native ranges

To compare crop climate suitability and crop area between the native and the expanded crop climatic ranges, we calculated the difference in mean climate suitability (and area) between native and current ranges. To aid in comparison between crops, we further divided this difference by the mean climate suitability (or cropland fraction) in the current niche. We tested for statistical significance of these standardized differences by randomly resampling 450 pixels of the native and current range 1000 times and calculated confidence of interval (α = 0.05) (Extended Data Figure 3). In doing so, we overcome the negative correlation between sample size and value.

### Pest data

We tested whether the presence of crop pests could explain the mismatches between crop area and climate suitability. However, since this hypothesis was not the main research question of this article, we restricted our analysis to the sunflower and its widespread pest *Sclerotinia sclerotiorm.* We focused on these two species because they have recently been the subject of a climatic niche study conducted at the global scale^10^, so that occurrence data of S. *sclerotiorm* were directly available. To test our hypothesis, we compared the mean of the mismatch between crop area and climate suitability (in absolute value) between pest-free and pest-presence areas. All analysis were conducted using R version 4.0.3.

## Data availability

The sources of all data used in this study are referenced in the Methods and all raw data are freely accessible at the URLs provided in Extended Data Table 2. The dataset used for the analyses is available from the corresponding author upon request.

## Code availability

The detailed script used for the analyses will be available online at the following URL: xxx

## Acknowledgements

LM’s work was funded by the French National Research Agency under the Programme “Investissements d’Avenir” under the reference ANR 17 MPGA 0004. LHR’s research on sunflower climate adaptation is funded by Genome Canada, Genome BC, and The Natural Sciences and Engineering Research Council of Canada. The Ministry of Economy and Competitivy of Spain (Grants CGL2014-56567-R and CGL2017-83855-R; Ministerio de Economía y Competitividad, Spain) fund RM research on crop’s wild progenitors. SP thanks the Bentham-Moxon Trust for funding a short stay at the University of British Columbia (BMT35-2017). C.K.K. was supported by grant no. 2019-67012-29733/project accession no. 1019405 from the USDA National Institute of Food and Agriculture. C.V. was supported by the European Research Council (ERC) Starting Grant Project “Ecophysiological and biophysical constraints on domestication in crop plants” (Grant ERC-StG-2014-639706-CONSTRAINTS). We thank Ian Ondo for his help with soil data, Emily Warschefsky and Navin Ramankutty for feedback on the manuscript. We thank Gilles Dauby and Thomas Couvreur for providing unpublished data from the Rainbio database. We also thank Nora Castañeda-Álvarez and Matija Obreza for extracting occurrence records from GENESYS and Dora A. L. Canhos and Sidnei De Souza for obtaining records from SpeciesLink database.

## Author contributions

L.M. led the data analysis and writing. L.M., S.P., D.R. and C.V. designed the experiment. F.B., C.K.K., R.M and C.P. contributed substantially to the crop wild progenitors’ analysis and writing. J.Y.B., S.P., D.R. and C.V. assisted with data analysis and writing. Z.M. and L.H.R. assisted with study design and writing.

## Additional information

Supplementary information, including Extended Data Tables and Extended Data Figures are available online.

### Extended Data Table Legends

**Extended Data Table 1. List of crop and associated wild progenitors.** N gives the number of occurrence (pixels) considered in the analysis.

**Extended Data Table 2. Source of data supporting findings**

**Extended Data Table 3. Outputs from the climate suitability models.**

MAT: mean annual temperature; TAP: total annual precipitation; %Var: Percentage of explained variance by each predictor; r^2^: adjusted coefficient of determination. Grey values show coefficients that are significant (see Methods)

**Extended Data Table 4: Outputs from the climate suitability models when accounting for climate seasonality.** MAT: mean annual temperature; TAP: total annual precipitation; %Var: Percentage of explained variance by each predictor; r^2^: adjusted coefficient. Grey values show coefficients that are significant (see Methods)

### Extended Data Figure Legends

**Extended Data Figure 1: Correlation between the fraction of available cropland devoted to each crop and climate suitability when accounting for temperature and precipitation seasonality.** Horizontal bars show confidence intervals (alpha = 0.05) computed from a random resampling procedure (see Methods). The correlation is considered significant if the confidence interval does not include 0. Black: non significant correlation; Blue: positive correlation; Red: negative correlation.

**Extended Data Figure 2**: Mismatch difference between *Sclerotinia sclerotiorum* free areas and areas where the pest occurs. *S. sclerotiorum* is a main sunflower’s pest worldwide. A positive mismatch score indicates that a low fraction of cultivated area is allocated to the crop while climate suitability is high. This mismatch is significantly lower in pixels with no pest (N = 129126) than with pests (N=66) (Wilcoxon test, W = 3586862, p-val = 0.026). Center line, median; box limits, upper and lower quartiles; whiskers, 1.5x interquartile range; points, outliers.

**Extended Data Figure 3**: Standardized differences in the fraction of cropland allocated to a crop (a) and climate suitability its yield (b) between native and current climatic ranges. Positive values indicate higher mean in the native range while negative values correspond to higher mean in the current range. Horizontal bars show confidence interval (alpha = 0.05) computed from 1000 random resamplings (see Methods). The difference is considered significant if the confidence interval does not include 0.

**Extended Data Figure 4**: Maps of climate suitability for crop production. Climate suitability is predicted for each crop by modelling the effects of mean annual temperature (MAT) and total annual precipitation (TAP) on crop yield (see Methods). Yellow colors represent zones of high climate suitability for crop production. Dark blue show zones of low climate suitability for crop production. Red polygons delineate the climate space occupies by crop wild progenitors.

**Extended Data Figure 5**: Maps of the fraction of available cropland allocated to each crop. Yellow colors represent zones where the crop covers large proportion of available cropland. Dark blue show zones where the crop occupies low fraction of available cropland. Red polygons delineate the climate space occupies by crop wild progenitors.

