## Supplementary Material for "Global mismatches between crop distributions and climate suitability"

### Extended Data – Tables

**Extended Data Table 1 - List of crop and associated wild progenitors.** N gives the number of occurrence (pixels) considered in the analysis.

| Crop | Crop wild progenitor | N crop | N progenitors |
|---|---|---|---|
| Barley | *Hordeum spontaneum* | 135617 | 85 |
| Cassava | *Manihot esculenta subsp. flabellifolia* | 100278 | 157 |
| Groundnut | *Arachis monticola*<br>*Arachis ipaensis*<br>*Arachis duranensis* | 119472 | 70 |
| Maize | *Zea mays subsp. parviglumis* | 176318 | 96 |
| Potato | *Solanum candolleanum* | 144651 | 331 |
| Rapeseed | *Brassica rapa subsp. oleifera*<br>*Brassica oleracea subsp. oleracea* | 107292 | 24 |
| Rice (asiatic) | *Oryza nivara*<br>*Oryza rufipogon* | 118110 | 1597 |
| Sorghum | *Sorghum bicolor subsp. verticilliflorum* | 138235 | 133 |
| Soybean | *Glycine soja* | 138663 | 309 |
| Sugarbeet | *Beta vulgaris subsp. maritima* | 52011 | 699 |
| Sunflower | *Helianthus annuus* | 129233 | 563 |
| Wheat | *Triticum urartu*<br>*Aegilops speltoides*<br>*Triticum dicoccoides*<br>*Aegilops tauschii*<br>*Triticum turgidum subsp. dicoccoides*<br>*Triticum monococcum subsp. boeoticum*<br>*Triticum monococcum subsp. aegilopoides*<br>*Triticum timopheevii subsp. armeniacum* | 162268 | 1174 |

**Extended Data Table 2 - Source of data supporting findings.**

| Layer | Name of the dataset | URL |
|---|---|---|
| *Agricultural dataset* | | |
| Crop yield | Harvested Area and Yield for 175 Crops year 2000 | http://www.earthstat.org/harvested-area-yield-175-crops/ |
| Harvested Area | Harvested Area and Yield for 175 Crops year 2001 | http://www.earthstat.org/harvested-area-yield-175-crops/ |
| Cropland Distribution | Cropland and Pasture Area in 2000 | http://www.earthstat.org/cropland-pasture-area-2000/ |
| *Agricultural management* | | |
| Fertilization | Nutrient Application for Major Crops | http://www.earthstat.org/nutrient-application-major-crops/ |
| Irrigation | The Global Map of Irrigation Areas (Version 4) | http://www.fao.org/aquastat/en/geospatial-information/global-maps-irrigated-areas/latest-version |
| *Climate data* | | |
| Mean annual temperature | CHELSA Bioclim | https://chelsa-climate.org/bioclim/ |
| Mean annual precipitation | CHELSA Bioclim | https://chelsa-climate.org/bioclim/ |
| Temperature seasonality | CHELSA Bioclim | https://chelsa-climate.org/bioclim/ |
| Precipitation seasonality | CHELSA Bioclim | https://chelsa-climate.org/bioclim/ |
| *Environmental and social economic data* | | |
| Soil pH | SoilGrids250m | https://data.isric.org/geonetwork/srv/fre/catalog.search#/search |
| Water capacity | SoilGrids250m | https://data.isric.org/geonetwork/srv/fre/catalog.search#/search |
| Soil organic content | SoilGrids250m | https://data.isric.org/geonetwork/srv/fre/catalog.search#/search |
| Slope | Slope 1KMmd GMTEDmd | http://www.earthenv.org/topography |
| Humand development index | HDI_1990_2015 | https://datadryad.org/stash/dataset/doi:10.5061/dryad.dk1j0 |
| Gross net Product | GDP_PPP_1990_2015_5arcmin | https://datadryad.org/stash/dataset/doi:10.5061/dryad.dk1j0 |
| *Crop wild progenitors occurrence records* | | |
| BIEN | Botanical Information and Ecology Network | https://bien.nceas.ucsb.edu/bien |
| GBIF | Global Biodiversity Information Facility | www.gbif.org |
| The Crop Trust | A global database for the distributions of crop wild relatives | www.cwrdiversity.org |
| BIOTIME | | http://biotime.st-andrews.ac.uk/ |
| GENESYS | Plant Genetic Resources for Food and Agriculture | https://www.genesys-pgr.org |
| RAINBIO | Sub-Saharan African vascular plants | http://rainbio.cesab.org/ |
| speciesLink | plant and animal occurrences in Brazil | http://splink.cria.org.br/ |

| Crop | MAT | | | TAP | | | Fertilisation | | | Irrigation | | | Soil pH | | Water Capacity | | Soil Organic Carbon | | Slope | | GDP | | HDI | | $r^2$ |
|---|---|---|---|---|---|---|---|---|---|---|---|---|---|---|---|---|---|---|---|---|---|---|---|---|---|
| | x | $x^2$ | %Var | x | $x^2$ | %Var | x | $x^2$ | %Var | x | $x^2$ | %Var | x | %Var | x | %Var | x | %Var | x | %Var | x | %Var | x | %Var | |
| **barley** | -0.02 | -0.05 | 6.06 | 0.17 | -0.06 | 16.32 | 0.19 | -0.01 | 54.75 | 0.03 | 0.00 | 0.41 | 0.09 | 6.26 | 0.04 | 2.20 | -0.03 | 1.64 | -0.03 | 2.24 | 0.00 | 0.01 | 0.08 | 10.12 | 0.36 |
| **cassava** | -0.09 | 0.05 | 8.27 | 0.13 | -0.05 | 2.75 | 0.21 | -0.01 | 19.29 | 0.23 | -0.03 | 11.00 | 0.09 | 1.94 | -0.02 | 0.19 | 0.00 | 0.00 | 0.01 | 0.06 | -0.01 | 0.07 | 0.27 | 56.44 | 0.30 |
| **groundnut** | 0.01 | 0.01 | 0.74 | 0.08 | -0.01 | 3.09 | 0.11 | 0.00 | 23.99 | 0.12 | -0.02 | 7.75 | 0.09 | 4.97 | 0.02 | 0.61 | 0.03 | 1.56 | 0.02 | 1.51 | 0.00 | 0.05 | 0.14 | 55.73 | 0.38 |
| **maize** | -0.18 | -0.07 | 11.04 | 0.19 | -0.05 | 4.67 | 0.24 | -0.02 | 31.32 | 0.06 | -0.01 | 0.79 | 0.19 | 9.19 | 0.02 | 0.21 | -0.03 | 0.41 | -0.03 | 0.97 | 0.00 | 0.00 | 0.26 | 41.40 | 0.58 |
| **potato** | 0.02 | -0.09 | 1.78 | 0.07 | -0.04 | 0.92 | 0.45 | -0.03 | 40.60 | 0.05 | 0.00 | 0.39 | 0.21 | 4.21 | 0.00 | 0.00 | -0.13 | 3.51 | -0.03 | 0.30 | 0.00 | 0.00 | 0.54 | 48.27 | 0.45 |
| **rapeseed** | 0.10 | -0.03 | 12.61 | 0.07 | -0.05 | 12.49 | 0.09 | 0.00 | 27.27 | -0.01 | 0.00 | 0.15 | 0.01 | 0.03 | 0.10 | 23.34 | -0.04 | 4.93 | -0.01 | 0.12 | 0.00 | 0.03 | 0.08 | 19.02 | 0.39 |
| **rice** | -0.08 | -0.01 | 2.76 | -0.01 | -0.03 | 2.90 | 0.31 | -0.02 | 74.68 | 0.08 | -0.01 | 1.44 | 0.09 | 2.78 | 0.01 | 0.15 | 0.02 | 0.47 | 0.02 | 0.58 | 0.00 | 0.00 | 0.11 | 14.23 | 0.51 |
| **sorghum** | -0.10 | -0.07 | 6.96 | 0.20 | -0.04 | 9.91 | 0.22 | -0.01 | 39.41 | 0.01 | -0.01 | 0.45 | 0.14 | 7.76 | 0.08 | 5.18 | -0.07 | 3.73 | 0.01 | 0.17 | 0.00 | 0.03 | 0.17 | 26.40 | 0.54 |
| **soybean** | 0.02 | -0.01 | 1.54 | 0.15 | -0.04 | 12.10 | 0.09 | -0.01 | 17.83 | -0.01 | 0.00 | 0.19 | 0.11 | 14.48 | 0.06 | 7.99 | -0.05 | 7.35 | 0.01 | 0.22 | -0.01 | 0.30 | 0.11 | 38.00 | 0.25 |
| **sugarbeet** | -0.22 | 0.08 | 1.58 | 0.37 | -0.06 | 2.08 | 1.00 | -0.13 | 48.76 | 0.18 | -0.01 | 1.09 | 0.35 | 2.59 | 0.04 | 0.07 | -0.30 | 3.71 | 0.14 | 1.06 | -0.03 | 0.05 | 0.86 | 39.00 | 0.48 |
| **sunflower** | 0.00 | 0.01 | 0.58 | 0.06 | -0.03 | 5.58 | 0.13 | -0.01 | 48.66 | 0.01 | 0.00 | 0.17 | -0.02 | 0.49 | 0.06 | 10.48 | -0.08 | 16.62 | -0.01 | 0.29 | 0.00 | 0.02 | 0.07 | 17.11 | 0.27 |
| **wheat** | 0.02 | -0.07 | 7.62 | 0.06 | -0.05 | 8.27 | 0.20 | -0.01 | 52.28 | -0.01 | 0.01 | 0.79 | 0.04 | 0.76 | 0.11 | 13.36 | -0.07 | 5.69 | -0.05 | 3.87 | 0.01 | 0.21 | 0.07 | 7.16 | 0.35 |

**Extended Data Table 3 - Outputs from the climate suitability models.** MAT: mean annual temperature; TAP: total annual precipitation; %Var: Percentage of explained variance by each predictor; $r^2$: adjusted coefficient of determination. Grey values show coefficients that are significant (see Methods).

| Crop | MAT | | | TAP | | | TempSeas | | PrecSeas | | Fertilisation | | | Irrigation | | | Soil pH | | Water Capacity | | Soil Organic Carbon | | Slope | | HDI | | $r^2$ |
|---|---|---|---|---|---|---|---|---|---|---|---|---|---|---|---|---|---|---|---|---|---|---|---|---|---|---|---|
| | x | $x^2$ | %Var | x | $x^2$ | %Var | x | %Var | x | %Var | x | $x^2$ | %Var | x | $x^2$ | %Var | x | %Var | x | %Var | x | %Var | x | %Var | x | %Var | |
| barley | -0.05 | -0.05 | 6.532 | 0.16 | -0.05 | 15.27 | -0.04 | 1.057 | 0.01 | 0.038 | 0.19 | -0.01 | 55.36 | 0.03 | 0 | 0.686 | 0.1 | 6.745 | 0.03 | 1.207 | -0.03 | 1.215 | -0.04 | 3.2 | 0.08 | 8.687 | 0.36 |
| cassava | 0.19 | 0.08 | 6.391 | 0.12 | -0.06 | 3.278 | 0.31 | 19.37 | -0.06 | 1.895 | 0.13 | -0.01 | 7.383 | 0.18 | -0.02 | 7.736 | 0.05 | 0.64 | 0.02 | 0.217 | -0.02 | 0.295 | 0.08 | 4.174 | 0.25 | 48.61 | 0.32 |
| groundnut | 0.03 | 0.02 | 1.452 | 0.07 | -0.01 | 3.134 | 0.01 | 0.103 | -0.02 | 1.08 | 0.11 | -0.01 | 25.29 | 0.12 | -0.02 | 9.317 | 0.09 | 5.859 | 0.02 | 0.54 | 0.02 | 1.337 | 0.03 | 2.274 | 0.13 | 49.56 | 0.39 |
| maize | -0.23 | -0.05 | 8.381 | 0.16 | -0.05 | 4.128 | -0.08 | 1.545 | -0.06 | 1.622 | 0.25 | -0.02 | 37.87 | 0.08 | -0.01 | 1.384 | 0.21 | 12.17 | 0 | 5E-06 | -0.03 | 0.566 | -0.05 | 1.608 | 0.24 | 30.71 | 0.59 |
| potato | -0.38 | -0.13 | 6.936 | 0.05 | -0.02 | 0.195 | -0.42 | 9.938 | 0.11 | 1.436 | 0.48 | -0.01 | 34.7 | 0.1 | -0.01 | 1.052 | 0.26 | 4.799 | -0.07 | 0.594 | -0.09 | 1.419 | -0.13 | 2.871 | 0.58 | 36.06 | 0.48 |
| rapeseed | -0.03 | -0.04 | 5.129 | 0.05 | -0.04 | 9.604 | -0.13 | 17.85 | 0.01 | 0.124 | 0.1 | -0.01 | 37.78 | 0 | -0.01 | 0.03 | 0.01 | 0.277 | 0.06 | 8.859 | -0.03 | 3.771 | -0.04 | 4.664 | 0.07 | 11.88 | 0.42 |
| rice | -0.07 | 0 | 1.774 | -0.03 | -0.03 | 3.589 | -0.03 | 0.335 | -0.05 | 2.173 | 0.33 | -0.02 | 77.08 | 0.08 | -0.01 | 1.736 | 0.11 | 3.68 | 0.01 | 0.041 | 0.01 | 0.125 | 0.02 | 0.365 | 0.09 | 9.102 | 0.51 |
| sorghum | -0.15 | -0.07 | 7.193 | 0.2 | -0.03 | 10.29 | -0.06 | 0.84 | 0.02 | 0.189 | 0.22 | -0.01 | 40.64 | 0.01 | -0.01 | 0.36 | 0.14 | 8.556 | 0.07 | 4.546 | -0.06 | 3.259 | 0 | 5E-04 | 0.18 | 24.09 | 0.54 |
| soybean | -0.13 | -0.03 | 7.229 | 0.12 | -0.03 | 5.839 | -0.18 | 17.96 | 0.01 | 0.062 | 0.13 | -0.01 | 22.66 | 0 | 0 | 0.007 | 0.12 | 13.57 | 0.02 | 0.759 | -0.03 | 2.26 | -0.03 | 1.972 | 0.12 | 27.34 | 0.29 |
| sugarbeet | -0.72 | 0.08 | 8.858 | 0.31 | -0.03 | 2.113 | -0.6 | 8.095 | -0.06 | 0.143 | 1 | -0.12 | 46.65 | 0.22 | -0.02 | 1.763 | 0.44 | 4.686 | -0.12 | 0.686 | -0.31 | 4.315 | -0.08 | 0.289 | 0.68 | 22.37 | 0.51 |
| sunflower | -0.07 | 0 | 2.874 | 0.06 | -0.03 | 4.603 | -0.07 | 4.123 | 0.02 | 1.419 | 0.13 | -0.01 | 46.84 | 0.02 | 0 | 0.483 | -0.01 | 0.162 | 0.06 | 7.986 | -0.07 | 12.23 | -0.02 | 1.731 | 0.08 | 17.55 | 0.28 |
| wheat | -0.1 | -0.07 | 8.763 | 0.05 | -0.04 | 5.302 | -0.13 | 8.171 | 0.01 | 0.125 | 0.21 | -0.01 | 52.67 | 0.01 | 0 | 1.041 | 0.05 | 1.38 | 0.08 | 5.079 | -0.05 | 3.282 | -0.08 | 8.018 | 0.08 | 6.03 | 0.36 |

**Extended Data Table 4 - Outputs from the climate suitability models when accounting for climate seasonality.** MAT: mean annual temperature; TAP: total annual precipitation; %Var: Percentage of explained variance by each predictor; $r^2$: adjusted coefficient. Grey values show coefficients that are significant (see Methods).

**Extended Data – Figures**

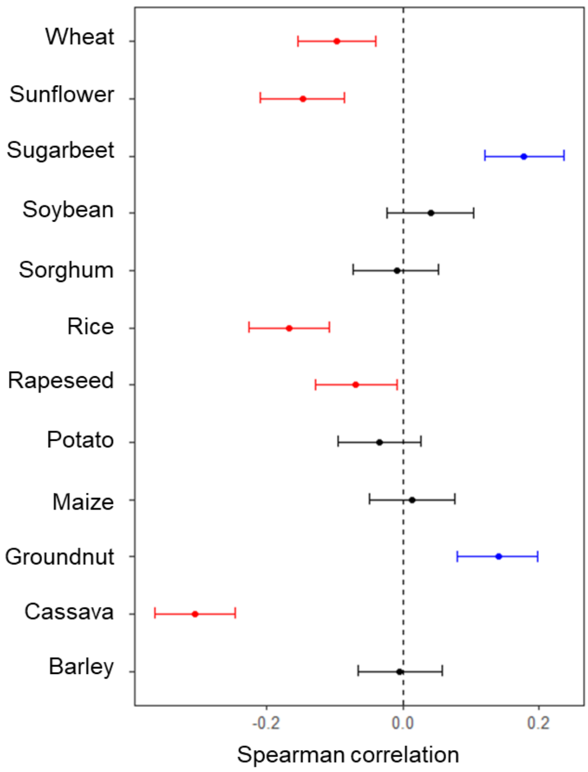


**Extended Data Figure 1 - Correlation between the fraction of available cropland devoted to each crop and climate suitability when accounting for temperature and precipitation seasonality.** Horizontal bars show confidence intervals (alpha = 0.05) computed from a random resampling procedure (see Methods). The correlation is considered significant if the confidence interval does not include 0. Black: non significant correlation; Blue: positive correlation; Red: negative correlation.

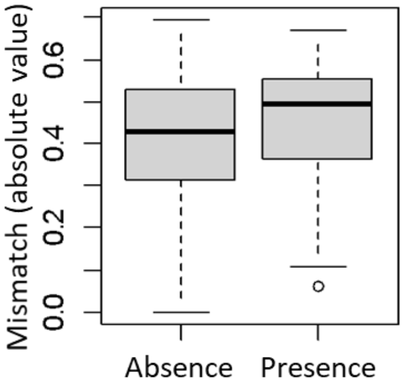


**Extended Data Figure 2: Mismatch difference between *Sclerotinia sclerotiorum* free areas and areas where the pest occurs.** *S. sclerotiorum* is a main sunflower's pest worldwide. A positive mismatch score indicates that a low fraction of cultivated area is allocated to the crop while climate suitability is high. This mismatch is significantly lower in pixels with no pest (N = 129126) than with pests (N=66) (Wilcoxon test, W = 3586862, p-val = 0.026). Center line, median; box limits, upper and lower quartiles; whiskers, 1.5x interquartile range; points, outliers.

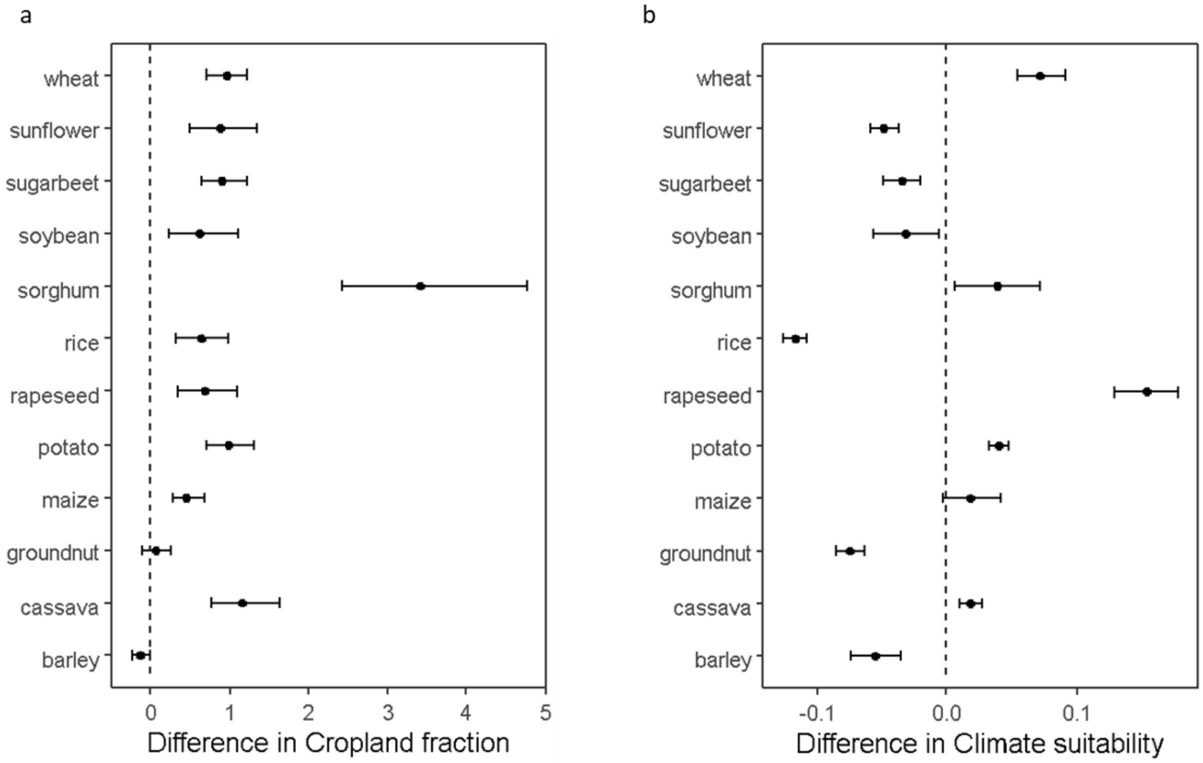


**Extended Data Figure 3 - Standardized differences in the fraction of cropland allocated to a crop (a) and climate suitability its yield (b) between native and current climatic ranges.** Positive values indicate higher mean in the native range while negative values correspond to higher mean in the current range. Horizontal bars show confidence interval (alpha = 0.05) computed from 1000 random resamplings (see Methods). The difference is considered significant if the confidence interval does not include 0.

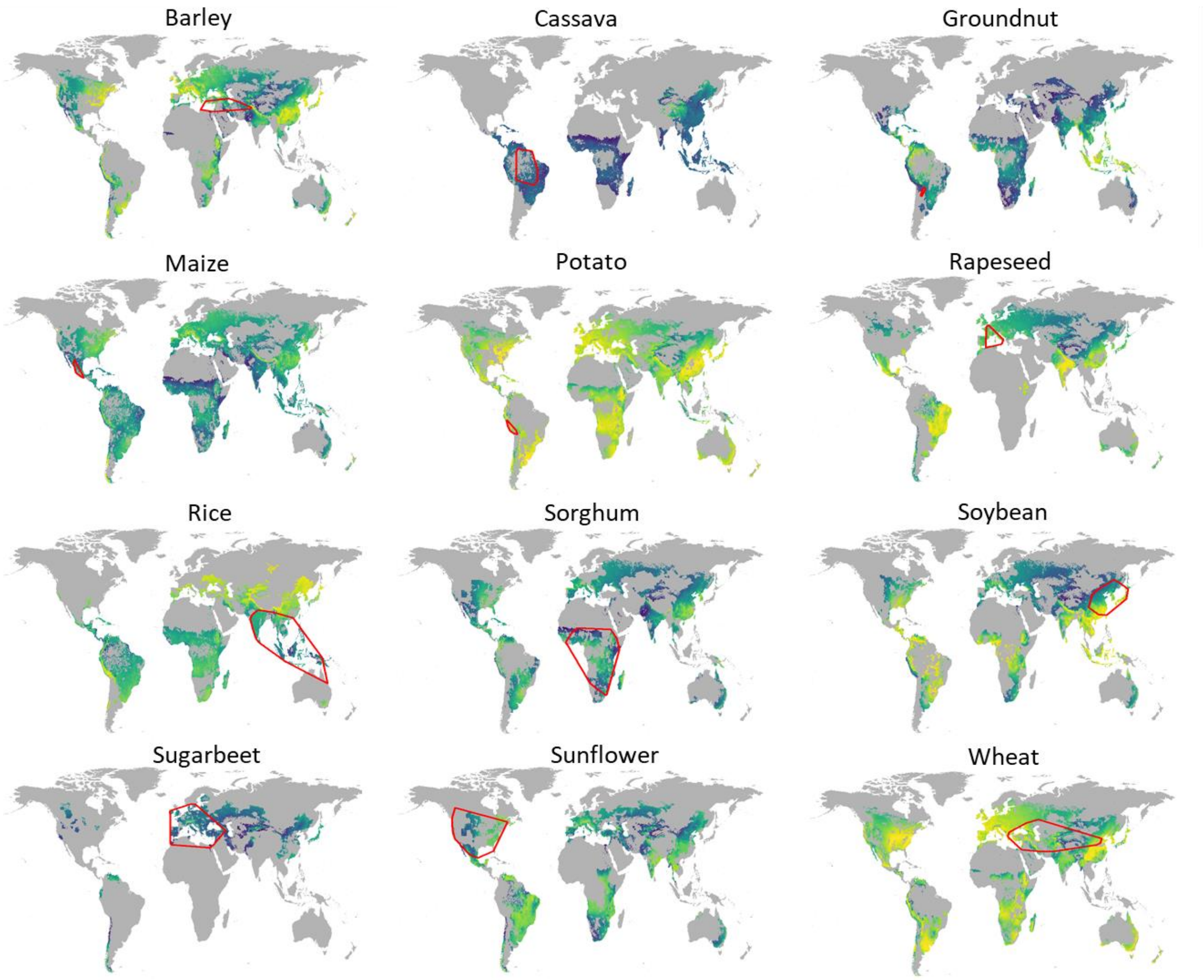


**Extended Data Figure 4 - Maps of climate suitability for crop production.** Climate suitability is predicted for each crop by modelling the effects of mean annual temperature (MAT) and total annual precipitation (TAP) on crop yield (see Methods). Yellow colors represent zones of high climate suitability for crop production. Dark blue show zones of low climate suitability for crop production. Red polygons delineate the climate space occupies by crop wild progenitors.

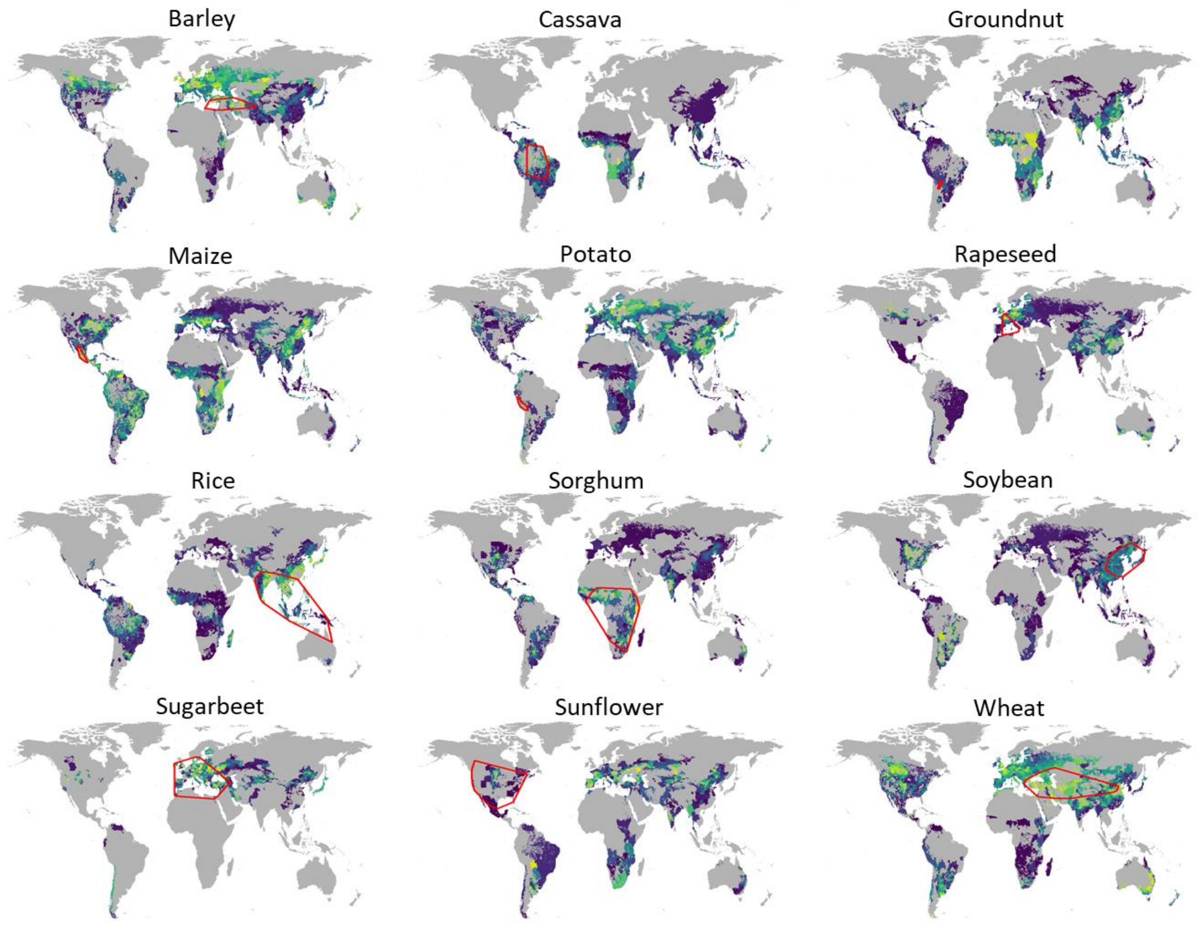


**Extended Data Figure 5 - Maps of the fraction of available cropland allocated to each crop.** Yellow colors represent zones where the crop covers large proportion of available cropland. Dark blue show zones where the crop occupies low fraction of available cropland. Red polygons delineate the climate space occupies by crop wild progenitors.
